# Reconstituted Mining Effluent Reduces Neuronal Proliferation in the Developing Brain and Slows Growth of Body and Facial Features in Wild-Caught Wood Frog Tadpoles

**DOI:** 10.1101/2020.01.29.924837

**Authors:** Hannah G. Sturgeon, Jeremy P. Kitchen, Lara I. Dahora, Sara E. Sweeten, Christopher K. Thompson

## Abstract

Mining, whether current or inactive, generally increases salt concentrations in catchment watersheds due to precipitation on and through exposed rock surfaces. Practices like mountaintop removal mining have exacerbated this issue, with measurements of salt concentrations in nearby catchment systems well above normal levels. Nevertheless, the impact of the ionic composition of mining effluent on aquatic animal health is not well understood. This is a particularly important issue in Appalachia because it is home to an enormous diversity of organisms, including a huge array of amphibians that live in streams that receive mining effluent from operating and abandoned mines. To investigate this issue, we examined the effects of reconstituted mining effluent on the development of wild-caught wood frog (Lithobates sylvaticus) tadpoles. We collected day-old fertilized eggs from a creek near Blacksburg, VA in early March, 2018 and raised them to hatch. Tadpoles were then assigned to either sulfate or chloride-based reconstituted mining effluent diluted to six different conductivities (100 μS/cm - 2,400 μS/cm). After 7 or 14 days of treatment, tadpoles were euthanized and fixed in paraformaldehyde. We imaged the heads and bodies of tadpoles for morphometric analysis before dissecting out brains and immunostaining them for phospho-histone H3, which labels dividing progenitor cells in the brain. We found that sulfate-based reconstituted mining effluent significantly lowered progenitor cell division at 1200 μS/cm at Day 7 and at 600 μS/cm at Day 14 relative to control. Chloride-based reconstituted mining effluent was less impactful, with no significant differences observed at Day 7 and significantly lowered progenitor cell division at 2400 μS/cm at Day 14. In addition, both treatments slowed growth of some head morphological features, including head size and interocular distance. Chloride treatment slowed growth of body length at Day 14 at 600 μS/cm, whereas sulfate-based reconstituted mining effluent had no effect on body length. These data show that sulfate-based mining effluent has a substantial impact on aspects of neural development, whereas chloride-based reconstituted mining effluent had less effect. In contrast, chloride-based reconstituted mining effluent had a much greater impact than sulfate on body morphology and growth. These experiments demonstrate that the chemical composition of salts in mining effluent can have divergent effects on the development of amphibians.

## Introduction

Since its inception, mining has been a major industry in the United States. Currently the mining industry provides over 500,000 jobs directly and indirectly more than a million jobs across other related fields (National Mining Association, 2018). While mining provides many jobs within largely rural communities, there is also potential for mines to have detrimental effects on the environment. In many cases, mining operations are unable to properly dispose of the waste rock and tailings produced by their mines, which then end up draining into nearby waterways (Salomons, 1995). When surveying a mining operation for overall external risk, government organizations often put a high priority on the potential dangers to the human population but sometimes neglect the potential effects on the welfare of the environment and wildlife present in that region (Veiga, Scoble, & McAllister, 2001). Some of this is due to a lack of data on the potential environmental effects of mining activities on wildlife. Elevated salt concentrations can be especially detrimental to aquatic life. A benchmark level of chronic conductivity of common salts was established by the EPA at 300 μS/cm, which signifies the level at which 95% of native species would still be alive (U.S. EPA, 2011).

Mining effluent is made up of a mixture of waste products that can include anything from the oil and grease used on machinery to heavy metals and salts (Tiwary, 2001; Sweeten et al., 2013). This unwanted material continuously emanates from both active and abandoned mines and ends up draining into nearby rivers and streams (Salomons, 1995). In addition, mines often have large areas of exposed earth and rock that are especially susceptible to erosion during rainfall (Tiwary, 2001), which can add to the debris load that flows into nearby waterways. Salts and heavy metals are the two components of mining effluent of the most environmental concern (Soucek & Kennedy, 2005; Goodfellow et al., 2000). Most often, studies investigating the presence of heavy metals in waterways affected by mining effluent report elevated concentrations of copper, zinc, nickel, and cadmium; (Dayeh, Lynn, & Bols, 2005; Mishra, Upadhyaya, Pandey, & Tripathi, 2008) but these findings can vary greatly depending on the geographical region studied. Chloride and sulfate salts are highly prevalent in mining effluent (Yeager-Armstead et al., 2013). While these salts are essential for the animal life within these streams, excessive levels of salts as a result of mining runoff create toxic conditions once they reach levels beyond those that animals are accustomed to (U.S. EPA, 2011). Mining effluent degrades water quality in part by lowering the pH and increasing the levels of dissolved solids (Tiwary, 2001). The approximate strength of dissolved salts can be estimated by quantifying the specific conductivity, which is the ability of a material to conduct an electric current and is often measured in microSiemens per centimeter (μS/cm) (U.S. EPA, 2011). In 1989, it was reported that approximately 19,300 km of rivers and streams across the United States had been degraded by mining effluent (Johnson & Hallberg, 2005). A more recent study estimated the changes made to the landscape of West Virginia between 1976 and 2005, and found that 5% of West Virginian land had been turned into surface mines, which in turn was affecting up to 56% of the rivers measured within the study area (Bernhardt et al., 2012). These measurements focus on a region of the nation in which mining is especially prevalent, but it is probable that similar increases were seen at other mining sites across the United States. In addition, it is likely that damage has increased in the time since these measurements were taken. Despite the ongoing research into the effects of mining effluent on the chemical makeup of natural waterways, the effects of these ions on amphibian well-being is not well understood.

To help fill in this gap, we investigated the impact of reconstituted mining effluent, with high concentrations of either chloride or sulfate ions, on morphological development and neurogenesis in the brains of wild-caught wood frog tadpoles.

## Methods

### Animals

We used wood frog (*Lithobates sylvaticus*) tadpoles. These animals were collected as day-old fertilized eggs from a creek near Blacksburg, VA, and then raised to hatch in the laboratory.

### Tadpole Collection

One day-old fertilized wood frog eggs were collected from a creek near Blacksburg, VA in March 2018. Prior to the date of collection, we visited the potential collection site daily to inspect the area for eggs. Eggs were collected on the first day that they were present, ensuring the age of the eggs. After collection, the tadpoles were raised to hatch in the laboratory in heat-controlled five gallon buckets.

### Treatment

Once tadpoles hatched, they were assigned to either a sulfate- or chloride-based reconstituted mining effluent group, at one of six conductivities ranging from 100 μS/cm - 2,400 μS/cm. The sulfate-based reconstituted mining effluent was created through combination of CaSO_4_, MgSO_4_, NaHCO_3_, KCl, and NaCl; and the chloride-based effluent through a similar procedure with a formula that was highly concentrated with chlorides, made up of CaCl_2_, MgCl_2_, NaHCO_3_, KCl, and NaCl. The tadpole groups were reared in solution for either seven or fourteen days.

### Sacrifice and Tissue Processing

Tadpoles were sacrificed on either Day 7 or Day 14 post-treatment (thus 14 or 21 days post fertilization), with an overdose of MS222 at 0.2%. Tadpoles were fixed in 4% phosphate-buffered paraformaldehyde (PFA) at 4°C. Tadpole brains were washed with PBS and then dissected into PBS-TX and placed in blocking buffer (2.5% norm al goat serum in PBS-TX) for 1 hour. Tadpole brains were incubated for 2 days in the primary antibody, anti-phospho-histone H3 (pH3) at 1:1000 (Sigma: H0412, made in rabbit) in blocking buffer at 4°C with gentle rotation. We then washed the brains three times in PBS-TX, and incubated them overnight in the secondary antibody, Alexa fluor 488 goat antirabbit IgG (1:400). Brains were washed in PBS-TX three times, then incubated in Sytox-O (Invitrogen: S34861), which labels cellular nuclei, in PBS at 1:500 once, washed in PBS and then mounted into custom-built wells on a slide with mounting media made from 50% Glycerol and 0.36% Urea in ddH_2_O.

### Imaging

Prior to brain dissection, the head and body of the tadpoles were imaged on a Nikon Model C-DSS115 microscope for later morphological analysis. After brains were dissected out and processed for immunostaining (above), they were imaged on a Leica SP8 confocal microscope. Images were taken at a 0.75 zoom on a 40X objective. Z-stack images with 100 steps were taken of Day 7 tadpole brains, while 150 steps were used for the Day 14 brains due to their larger size. Each hemisphere was imaged separately. After the confocal brain images were analyzed, the colors were changed to magenta and green so that they could be understood properly by all audiences.

### Analysis

Morphological analysis was done in ImageJ. Length measurements were taken using the segmented line tool for body length (from snout to tail tip) and interocular distance (from outer part of left eye to outer part of right eye). The area of the head was taken using the polygon selection tool tracing the outer circumference of the head from the dorsal perspective. Neuronal analysis was done in Imaris. 3D models were created from confocal stacks using the surface tool to section off the region of interest from the 3D image. The volume of the 3D region of interest was measured using the statistics tool, and the density of pH3+ cells within the region of interest was quantified by manual counting and dividing by the volume of the region of interest.

### Statistics

GraphPad Prism was used for statistical analysis. We ran one-way ANOVA tests with a Dunnett’s multiple comparisons test, comparing the mean of each column with the mean of the control column, for each of the treatment groups (100 μS/cm – 2400 μS/cm). Scatter dot plots were created in GraphPad.

## Results

### Sulfate-based reconstituted mining effluent lowers progenitor cell division

To test the effects of sulfate-based reconstituted mining effluent on cell proliferation, we treated tadpoles with a range of conductivity of sulfate-based reconstituted mining effluent. Tadpoles were sacrificed at either Day 7 or Day 14, and were stained with pH3 and Sytox-O. The tadpoles sacrificed at Day 7 showed a significant reduction in cell proliferation at the conductivity level of 1200 μS/cm compared to the tadpoles treated with a baseline level of 100 μS/cm. The tadpoles sacrificed at Day 14 showed a significant reduction in cell proliferation at the conductivity level of 600 μS/cm compared to baseline.

### Chloride-based reconstituted mining effluent lowers progenitor cell division at Day 14

To test the effects of chloride-based reconstituted mining effluent on cell proliferation, we treated tadpoles with a range of conductivity of chloride-based reconstituted mining effluent. Tadpoles were sacrificed at either Day 7 or Day 14, and were stained with pH3 and Sytox-O. The tadpoles sacrificed at Day 7 did not show a reduction in cell proliferation at any elevated level of effluent conductivity when compared to the baseline level of 100 μS/cm. The tadpoles sacrificed at Day 14 showed a significant reduction in cell proliferation at the conductivity level of 2400 μS/cm compared to baseline.

### Chloride-based reconstituted mining effluent slows growth of head size and interocular distance at Day 14

Images of tadpole heads were used to analyze two features, head size and interocular distance. These images allowed us to examine the effects of chloride-based mining effluent at a range of conductivity on the growth of morphological features. The tadpoles sacrificed at Day 7 did not show a significant reduction in the growth of head size or interocular distance compared to baseline. The tadpoles sacrificed at Day 14 showed a significant reduction in the growth of both head size and interocular distance at the conductivity level of 1800 μS/cm relative to baseline.

### Sulfate-based reconstituted mining effluent slows growth of head size and interocular distance

Images of tadpole heads were used to analyze two features, head size and interocular distance. These images allowed us to examine the effects of sulfate-based mining effluent at a range of conductivity on the growth of morphological features. The tadpoles sacrificed at Day 7 showed a significant reduction in the growth of both head size and interocular distance at the conductivity level of 1800 μS/cm relative to baseline. The tadpoles sacrificed at Day 14 showed a significant reduction in the growth of head size at 1800 μS/cm relative to baseline, but no significant effects on interocular distance were seen.

### Chloride-based reconstituted mining effluent slows growth of full animal length at Day 14

Images of tadpole bodies were used to analyze the effects of chloride-based mining effluent at a range of conductivity on the growth of the full animal. Tadpoles sacrificed at Day 7 did not show a significant reduction in the length of the full animal relative to baseline. Tadpoles sacrificed at Day 14 showed a significant reduction in the length of the full animal at the conductivity level of 600 μS/cm relative to baseline.

### Sulfate-based reconstituted mining effluent has no significant effects on growth of the full animal

Images of tadpole bodies were used to analyze the effects of sulfate-based mining effluent at a range of conductivity on the growth of the full animal. Tadpoles sacrificed at both Day 7 and Day 14 showed no significant reductions in the length of the full animal relative to baseline.

## Discussion

In this study we investigated the effects of chloride-based and sulfate-based reconstituted mining effluent on the development of both the brain and morphological features of wild-caught wood frog tadpoles. Sulfate-based mining effluent significantly lowered progenitor cell division in the optic tectum at 1200 μS/cm at Day 7 and at 600 μS/cm at Day 14 relative to control, and chloride-based mining effluent significantly lowered progenitor cell division in the optic tectum at 2400 μS/cm at Day 14. These data suggest that sulfate-based mining effluent has a greater effect than chloride-based mining effluent on measures related to brain development. In contrast, we found that chloride-based mining effluent had more extreme effects on measures of morphological growth, such as the growth of tadpole head and body and interocular distance, when compared to sulfate-based mining effluent.

The fact that sulfate-based treatment had a more pronounced impact on proliferation than chloride-based treatment may be due to the role of sulfates in coordinating developmental critical periods. Perineuronal nets (PNN) are an important determinant of neuronal development. The appearance of PNNs typically marks the end of developmental critical periods, leading to stabilization of neural circuits (Kwok, Foscarin, & Fawcett, 2014). PNNs are made up of four classes of extracellular matrix molecules, one being chondroitin sulfate proteoglycans (CSPG) (Sorg et al., 2016). CSPGs are composed of a core protein and a chondroitin-sulfate-glycosaminoglycan chain (Miyata & Kitagawa, 2017). The sulfation profile in the glycosaminoglycan chain regulates PNN formation (Fawcett, Oohashi, & Pizzorusso, 2019). The glycosaminoglycan chains are most often sulfated in the 4 (C4S) or 6 (C6S) positions (Foscarin, et al., 2017). In early development the glycosaminoglycan chains are sulfated in the C6S position, and the prevalence of C6S sulfation decreases throughout development, leading to a final decline at the end of the critical period. During this time, the prevalence of C4S sulfation gradually increases, and takes predominance after the closure of the critical period (Foscarin, et al., 2017). Sulfate-based reconstituted mining effluent may lead to disturbance of the sulfation position within the glycosaminoglycan chains, potentially altering the critical period and stabilization of developing neural circuits. Given the decrease in proliferation observed here, we predict that sulfate-based treatment may increase C4S sulfation, thereby stabilizing PNNs too soon, and inducing early termination of the critical period.

Superficially one may predict that chloride-based mining effluent treatment would have greater effects on brain development than sulfate-based treatment, due to the fact that chloride is a critical ion for normal brain function (Elorza-Vidal, Gaitán-Peñas, & Estévez, 2019). The fact that chloride-based treatment only affected the brain at 14 days and only at the highest concentration demonstrates that the tadpole nervous system is particularly resistant to high levels of chloride exposure. Due to its omnipresence within the brain, it is possible that neurogenesis may have been less impaired by chloride-based treatment due to homeostatic mechanisms already in place to balance fluctuations in chloride levels. There may still be other changes in the brains of chloride-based treatment tadpoles, however; in particular we predict changes in the expression of molecular mechanisms that control chloride homeostasis.

While sulfate-based treatment more significantly affected neuronal proliferation, chloride-based treatment had greater effects on body measurements. This could be due to a variety of effects resulting from the high chloride levels, but we suspect it may be due to a decrease in water retention. Natterjack toad (*Bufo calamita*) tadpoles are known to regulate their water retention based on the environmental osmolality. Once osmolality reaches 90 mOsm, the Natterjack toad tadpoles begin reducing their water retention (Gomez-Mestre et al., 2004). It is likely that wood frog tadpoles would have similar capabilities, which may have led to the reduction in size seen as a result of chloride-based treatment.

In addition to the toxic effects these high levels of ion concentrations can cause by themselves, these levels can also create inhospitable conditions characterized by increased water hardness and low pH, which lead to additional toxic effects in amphibians. An increase in water hardness is thought to make a water source undrinkable for the human population (Tiwary, 2001). Water hardness is typically due to high levels of calcium carbonate, and therefore is often referred to in terms of “mg/L, as CaCO_3_”, but this term can also describe elevated quantities of other compounds composed of ions such as chloride and sulfate, as would be present in mining effluent (David J. Soucek et al., 2011). The potential toxicity of elevated water hardness to humans suggests that it could be equally or more significantly dangerous to wood frog tadpoles. Acid mine drainage (AMD), a condition in which sulfide-bearing minerals interact with both oxygen and water creating highly acidic runoff, has a huge potential to deposit sulfate ions into waterways (Akcil & Koldas, 2006). This process can be intensified by the introduction of a bacteria, such as thiobacillus bacteria, which can lead to an even greater increase in acidification (Tiwary, 2001). There are some cases in which this event has occurred naturally, but the majority of these incidences arise as a consequence of mining operations (Gray, 1997). AMD is a complex toxicological threat, in that it is capable of causing a wide range of environmental complications. In rivers and streams, the most hazardous effects to animal life are the decreases in pH of the entire water system and the metal toxicity that occurs as a result of AMD (Gray, 1997). These effects can sometimes be rectified by the addition of lime to the runoff, neutralizing the acid that has formed (Akcil & Koldas, 2006). Through the use of sulfate ions in our mining effluent treatment, we may have created an acidic environment for the tadpoles that mirrors the conditions seen in a river effected by AMD.

The significant effects of these sulfate and chloride-based treatments on brain and morphological development in wood frog tadpoles supports the idea that potential behavior changes could be present in these tadpoles as well. One study used the *Oncorhynchus mykiss*, or rainbow trout, as a biosensitive animal model to test the effects of mining effluent on two measures of behavior, locomotion and ventilation (Gerhardt, 1998). They created a reconstituted mining effluent treatment with high concentrations of both chloride and sulfate and treated the fish over a period of four days. They saw immediate behavioral effects in the fish, characterized by decreases in locomotor activity and increased time spent on ventilation (Gerhardt, 1998). It is possible that similar effects would be seen in wood frog tadpoles subjected to high levels of chloride and sulfate, and future experiments should address the impact of reconstituted mining effluent on tadpole and frog behavior.

Our data suggest that mining effluent has a significant impact on amphibian health, and effluent that is high in sulfate ions is especially detrimental to anuran brain development. Due to these and other deleterious effects of sulfate, methods for the removal of sulfate ions from streams and rivers may be vital to the well-being of aquatic life. Filtration is one of the most commonly used methods of sulfate removal, and is most often done through the addition of polyelectrolytes that will induce precipitation of the sulfate ions, allowing for their removal (Bowell, 2004). Furthermore, effluent dominated with chloride impacts aspects of development, although the long-term effects of chloride exposure early on in development is not known. Ultimately, it is clear that the type of mining effluent can have differential impacts on aspects of development, and that attention must be paid to specific organ systems.

---

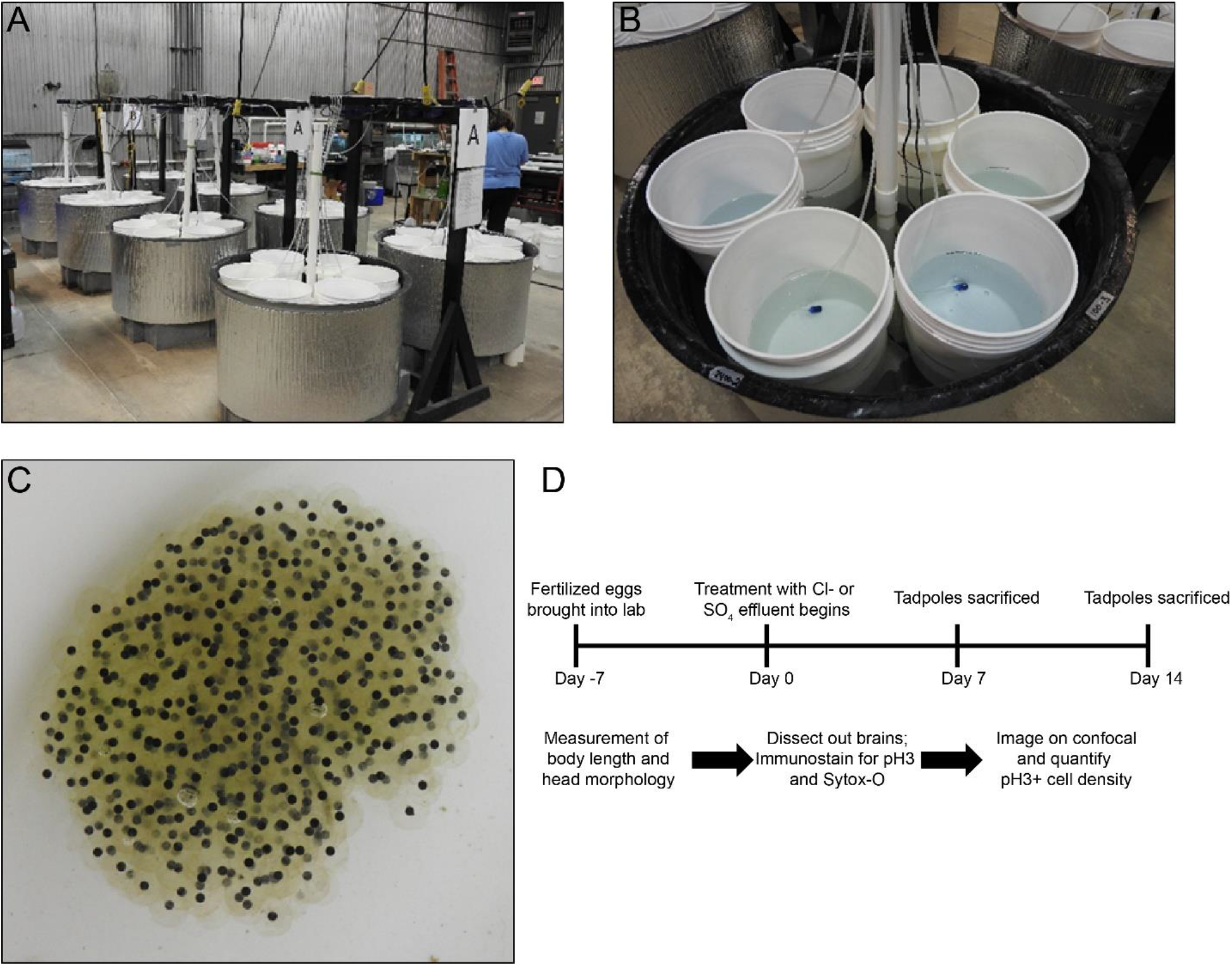
<u>Experimental set-up and timeline</u> A) Illustration of facility used in this experiment. Tadpoles were raised in a temperature controlled environment. 50-gallon water baths with six 5-gallon buckets in each water bath. B) Image of one water bath with six 5-gallon buckets filled with treatment. Treatment type and concentration was randomly distributed across four 50 gallon water baths (24 treatments in total). Each bucket contained 10 liters of treatment solution and an air stone. C) Wood frog eggs collected from Price’s Fork, Virginia on February 28, 2018. D) Timeline illustrating treatment regimen and experimental procedures.

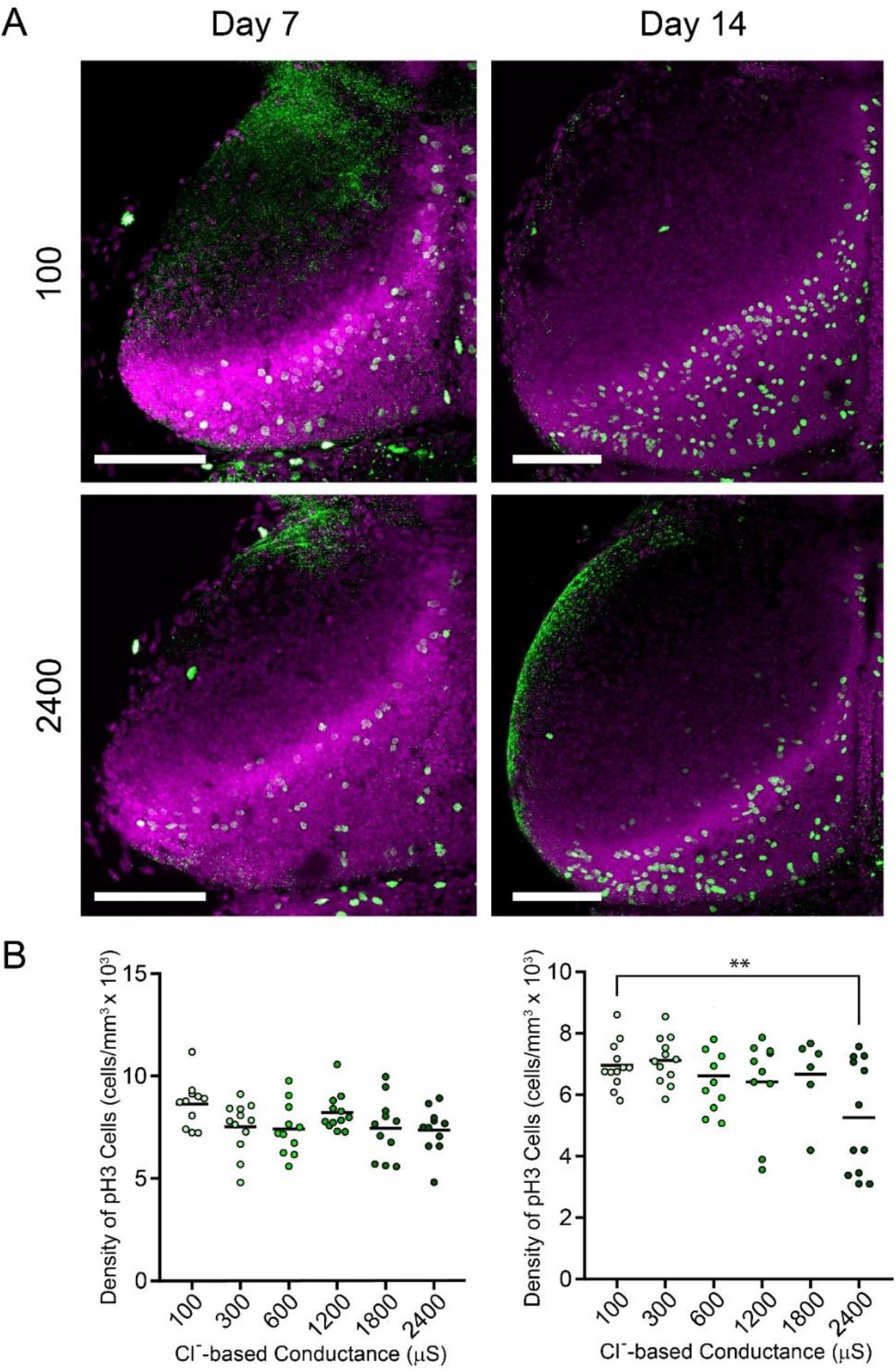
Chloride-based reconstituted mining effluent significantly lowered progenitor cell division at 2400 μS/cm relative to control at Day 14. A) Z-projections of confocal stacks of example optic tecta at 100 μS/cm and 2400 μS/cm from Day 7 and Day 14. B) Scatter plots showing proliferating cells for tadpoles sacrificed on Day 7 and Day 14. Scale bars represent 200 μm. ** = p< 0.01

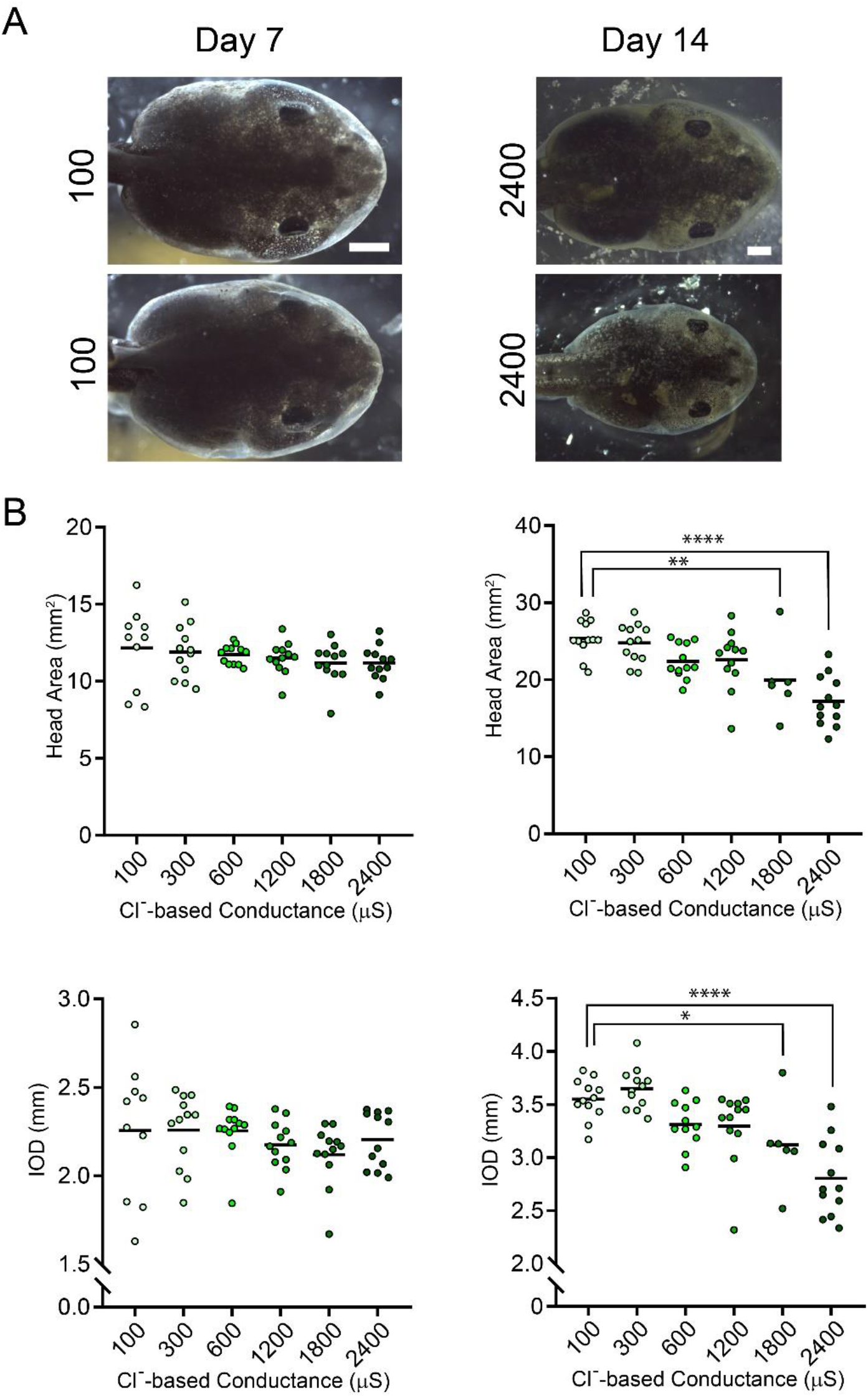
Chloride-based reconstituted mining effluent had no significant effect on morphological features relative to control at Day 7, and significantly lowered head size and interocular distance at 1800 μS/cm relative to control at Day 14. A) Example images of tadpole heads at Day 7 and Day 14, at both 100 μS/cm and 2400 μS/cm exhibiting head size and interocular distance. B) Scatter plots showing the effects of chloride-based mining effluent on the size of morphological features such as head size and interocular distance. Scale bars represent 1 mm. * = p<0.05, ** = p< 0.01, **** = p< 0.0001

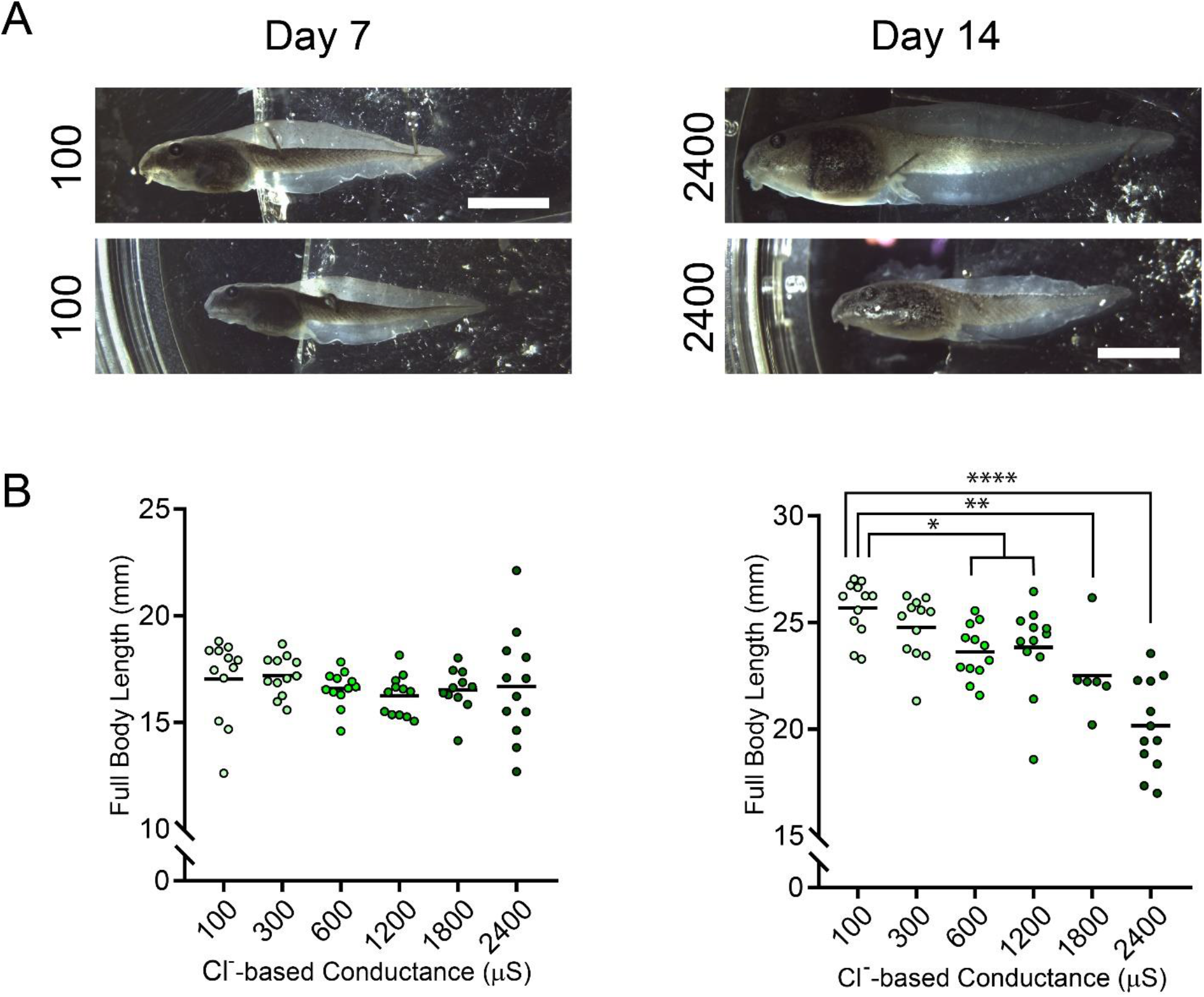
Chloride-based reconstituted mining effluent had no significant effect on full animal length at Day 7 relative to control, and significantly lowered full animal length at 600 μS/cm relative to control at Day 14. A) Example images of tadpole bodies at Day 7 and Day 14, at both 100 μS/cm and 2400 μS/cm exhibiting full body length. B) Scatter plots showing the effects of chloride-based mining effluent on the size of full animal length. Scale bars represent 5 mm. * = p<0.05, ** = p< 0.01, **** = p< 0.0001

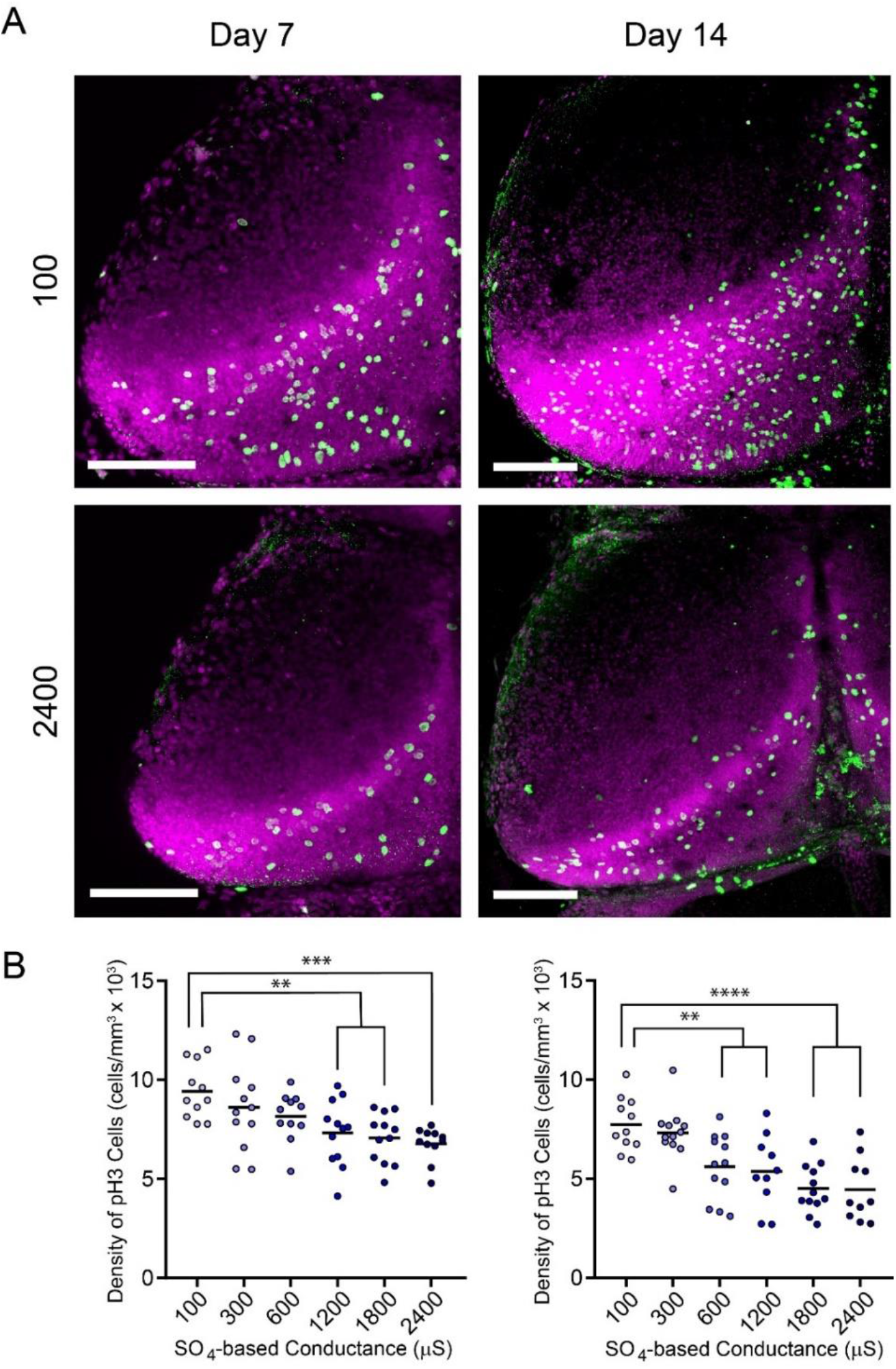
Sulfate-based reconstituted mining effluent significantly lowered progenitor cell division at 1200 μS/cm at Day 7, and at 600 μS/cm at Day 14 relative to control. A) Z-projections of confocal stacks of example optic tecta at 100 μS/cm and 2400 μS/cm from Day 7 and Day 14. B) Scatter plots showing proliferating cells for tadpoles sacrificed on Day 7 and Day 14. Scale bars represent 200 μm. ** = p< 0.01, *** = p< 0.001, **** = p< 0.0001

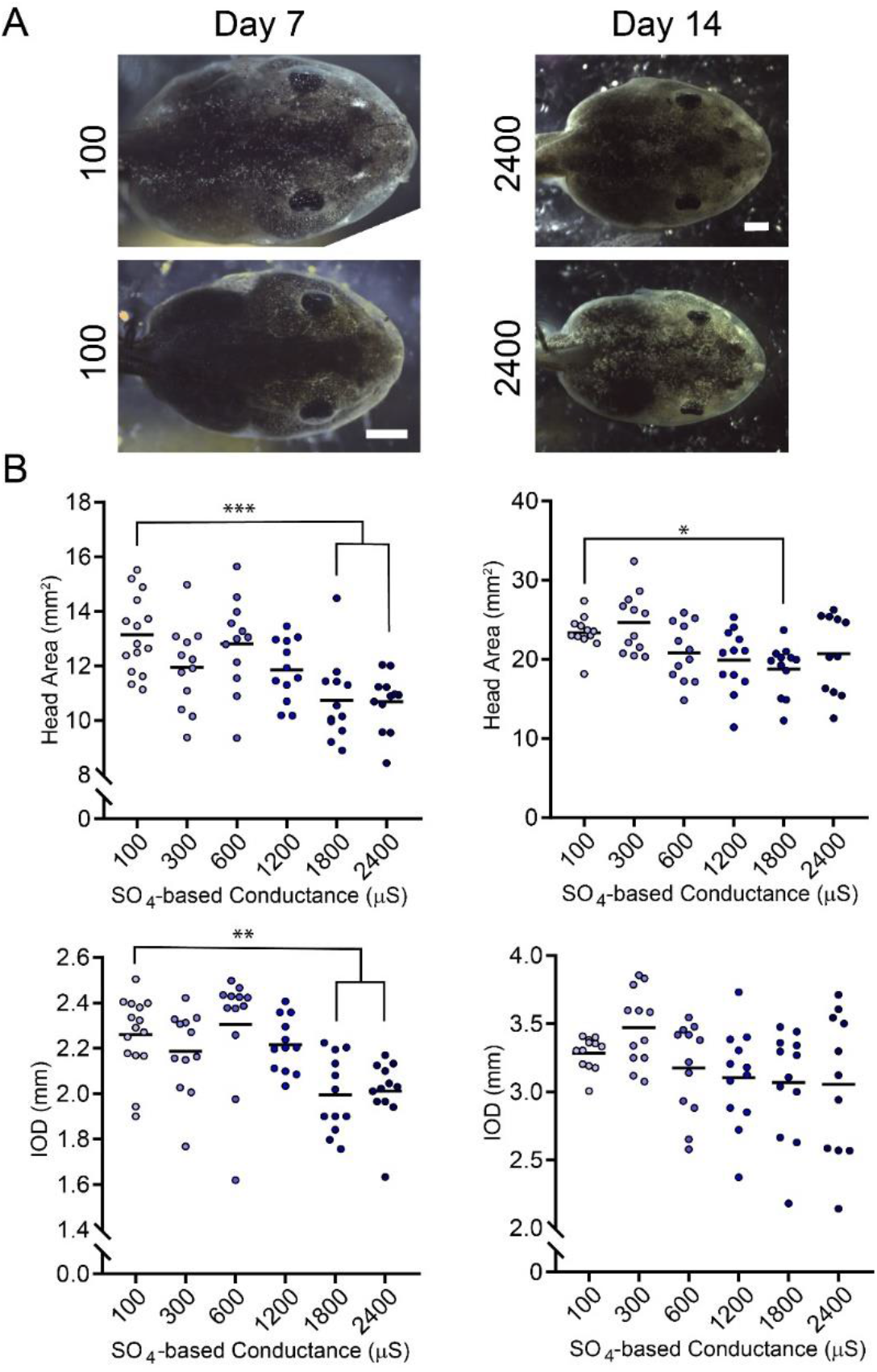
Sulfate-based reconstituted mining effluent significantly lowered head size at 1800 μS/cm relative to control. Sulfate-based reconstituted mining effluent significantly lowered interocular distance at 1800 μS/cm relative to control at Day 7 and had no significant effects on interocular distance relative to control at Day 14. A) Example images of tadpole heads at Day 7 and Day 14, at both 100 μS/cm and 2400 μS/cm exhibiting head size and interocular distance. B) Scatter plots showing the effects of sulfate-based mining effluent on the size of morphological features such as head size and interocular distance. Scale bars represent 1 mm. * = p<0.05, ** = p< 0.01, *** = p< 0.001

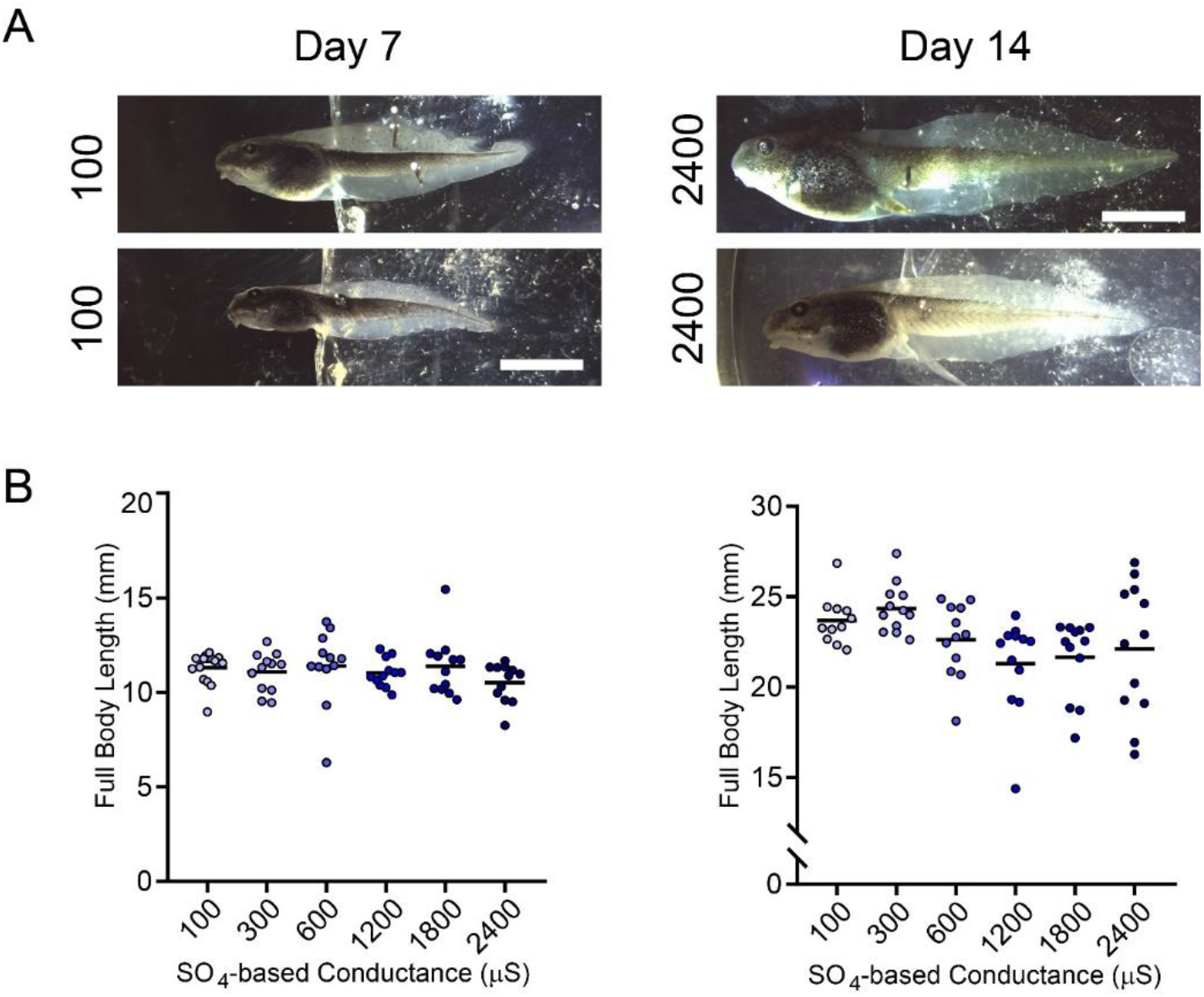
Sulfate-based reconstituted mining effluent had no significant effects on full animal length relative to control. A) Example images of tadpole bodies at Day 7 and Day 14, at both 100 μS/cm and 2400 μS/cm exhibiting full body length. B) Scatter plots showing the effects of sulfate-based mining effluent on the size of full animal length. Scale bars represent 5 mm.

## References

Akcil, A., & Koldas, S. (2006). Acid Mine Drainage (AMD): causes, treatment and case studies. Journal of Cleaner Production, 14(12-13 SPEC. ISS.), 1139–1145. https://doi.org/10.1016/j.jclepro.2004.09.006

National Mining Association. (2018). The Economic Contributions of U.S. Mining.

Bernhardt, E. S., Lutz, B. D., King, R. S., Fay, J. P., Carter, C. E., Helton, A. M., Campagne, D., & Amos, J. (2012). How Many Mountains Can We Mine? Assessing the Regional Degradation of Central Appalachian Rivers by Surface Coal Mining. Environmental Science & Technology, 46(15), 8115–8122. https://doi.org/10.1021/es301144q

Bowell, R. J. (2004). A review of sulphate removal options for mine waters. In Proceedings International Mine Water Association Symposium (Vol. 2, pp. 75–91). Newcastle.

Dayeh, V. R., Lynn, D. H., & Bols, N. C. (2005). Cytotoxicity of metals common in mining effluent to rainbow trout cell lines and to the ciliated protozoan, Tetrahymena thermophila. Toxicology in Vitro, 19(3), 399–410. https://doi.org/10.1016/j.tiv.2004.12.001

Elorza-Vidal, X., Gaitán-Peñas, H., & Estévez, R. (2019). Chloride channels in astrocytes: Structure, roles in brain homeostasis and implications in disease. International Journal of Molecular Sciences, 20(5). https://doi.org/10.3390/ijms20051034

Fawcett, J. W., Oohashi, T., & Pizzorusso, T. (2019). The roles of perineuronal nets and the perinodal extracellular matrix in neuronal function. Nature Reviews Neuroscience. https://doi.org/10.1038/s41583-019-0196-3

Foscarin, S., Raha-Chowdhury, R., Fawcett, J. W., & Kwok, J. C. F. (2017). Brain ageing changes proteoglycan sulfation, rendering perineuronal nets more inhibitory. Aging, 9(6), 1607–1622. https://doi.org/10.18632/aging.101256

Gerhardt, A. (1998). Whole Effluent Toxicity Testing with Oncorhynchus mykiss (Walbaum 1792): Survival and Behavioral Responses to a Dilution Series of a Mining Effluent in South Africa. Archives of Environmental Contamination and Toxicology, 35(2), 309–316. https://doi.org/10.1155/2019/3746964

Gomez-Mestre, I., Tejedo, M., Ramayo, E., Estepa, J., & Luisa, M. (2004). Developmental Alterations and Osmoregulatory Physiology of a Larval Anuran under Osmotic Stress. Physiological and Biochemical Zoology, 77(2), 267–274.

Goodfellow, W. L., Ausley, L. W., Burton, D. T., Denton, D. L., Dorn, P. B., Grothe, D. R., Heber, M. A., Norberg-King, T. J., & Rodgers, J. H. (2000). Major ion toxicity in effluents: A review with permitting recommendations. Environmental Toxicology and Chemistry, 19(1), 175–182. https://doi.org/10.1002/etc.5620190121

Gray, N. F. (1997). Environmental impact and remediation of acid mine drainage: a management problem. Environmental Geology, 30(2).

Johnson, D. B., & Hallberg, K. B. (2005). Acid mine drainage remediation options: A review. Science of the Total Environment, 338(1–2), 3–14. https://doi.org/10.1016/j.scitotenv.2004.09.002

Kwok, J. C. F., Foscarin, S., & Fawcett, J. W. (2014). Perineuronal nets: A special structure in the central nervous system extracellular matrix. In Extracellular Matrix (pp. 23–32). Springer New York. https://doi.org/10.1007/978-1-4939-2083-9_3

Mishra, V. K., Upadhyaya, A. R., Pandey, S. K., & Tripathi, B. D. (2008). Heavy metal pollution induced due to coal mining effluent on surrounding aquatic ecosystem and its management through naturally occurring aquatic macrophytes. Bioresource Technology, 99(5), 930–936. https://doi.org/10.1016/j.biortech.2007.03.010

Miyata, S., & Kitagawa, H. (2017, October 1). Formation and remodeling of the brain extracellular matrix in neural plasticity: Roles of chondroitin sulfate and hyaluronan. Biochimica et Biophysica Acta - General Subjects. Elsevier B.V. https://doi.org/10.1016/j.bbagen.2017.06.010

Salomons, W. (1995). Environmental impact of metals derived from mining activities: Processes, predictions, prevention. Journal of Geochemical Exploration, 52(1–2), 5–23. https://doi.org/10.1016/0375-6742(94)00039-E

Sorg, B. A., Berretta, S., Blacktop, J. M., Fawcett, J. W., Kitagawa, H., Kwok, J. C. F., & Miquel, M. (2016). Casting a wide net: Role of perineuronal nets in neural plasticity. Journal of Neuroscience, 36(45), 11459–11468. https://doi.org/10.1523/JNEUROSCI.2351-16.2016

Soucek, David J., Linton, T. K., Tarr, C. D., Dickinson, A., Wickramanayake, N., Delos, C. G., & Cruz, L. A. (2011). Influence of water hardness and sulfate on the acute toxicity of chloride to sensitive freshwater invertebrates. Environmental Toxicology and Chemistry, 30(4), 930–938. https://doi.org/10.1002/etc.454

Soucek, David John, & Kennedy, A. J. (2005). EFFECTS OF HARDNESS, CHLORIDE, AND ACCLIMATION ON THE ACUTE TOXICITY OF SULFATE TO FRESHWATER INVERTEBRATES. Environmental Toxicology and Chemistry, 24(5), 1204. https://doi.org/10.1897/04-142.1

Sweeten, S., Sweeten, J., Craynon, J., Ford, W. M., & Schoenholtz, S. (2013). Evaluation of Current Measures of Aquatic Biological Integrity in the Central Appalachian Coalfields: Efficacy and Implications. In J. Craynon (Ed.), Environmental Considerations in Energy Production (pp. 381–394). Englewood, Colorado, USA: Society for Mining, Metallurgy, and Exploration, Inc.

Tiwary, R. K. (2001). Environmental impact of coal mining on water regime and its management. Water, Air, and Soil Pollution, 132(1–2), 185–199. https://doi.org/10.1023/A:1012083519667

U.S. EPA (Environmental Protection Agency). 2011. A Field-Based Aquatic Life Benchmark for Conductivity in Central Appalachian Streams. Office of Research and Development, National Center for Environmental Assessment, Washington, DC. EPA/600/R-10/023F.

Veiga, M. M., Scoble, M., & McAllister, M. L. (2001). Mining with communities. Natural Resources Forum, 25(3), 191–202. https://doi.org/10.1111/j.1477-8947.2001.tb00761.x

Yeager-Armstead, M., Wilson, M., Keller, L., Kinney, J., McGill, K., & Snyder, E. (2013). Methods for evaluating the effects of a simulated mine effluent with elevated ionic concentration to field collected benthic macroinvertebrates.

